# Hierarchical Ensembles of Intrinsically Disordered Proteins at Atomic Resolution in Molecular Dynamics Simulations

**DOI:** 10.1101/731133

**Authors:** Lisa M. Pietrek, Lukas S. Stelzl, Gerhard Hummer

## Abstract

Intrinsically disordered proteins (IDPs) constitute a large fraction of the human proteome and are critical in the regulation of cellular processes. A detailed understanding of the conformational dynamics of IDPs could help to elucidate their roles in health and disease. However the inherent flexibility of IDPs makes structural studies and their interpretation challenging. Molecular dynamics (MD) simulations could address this challenge in principle, but inaccuracies in the simulation models and the need for long simulations have stymied progress. To overcome these limitations, we adopt an hierarchical approach that builds on the “flexible meccano” model of Bernadó et al. (J. Am. Chem. Soc. 2005, 127, 17968-17969). First, we exhaustively sample small IDP fragments in all-atom simulations to capture local structure. Then, we assemble the fragments into full-length IDPs to explore the stereochemically possible global structures of IDPs. The resulting ensembles of three-dimensional structures of full-length IDPs are highly diverse, much more so than in standard MD simulation. For the paradigmatic IDP *α*-synuclein, our ensemble captures both local structure, as probed by nuclear magnetic resonance (NMR) spectroscopy, and its overall dimension, as obtained from small-angle X-ray scattering (SAXS) in solution. By generating representative and meaningful starting ensembles, we can begin to exploit the massive parallelism afforded by current and future high-performance computing resources for atomic-resolution characterization of IDPs.

## INTRODUCTION

Proteins with intrinsically disordered regions constitute a large fraction of the human proteome.^1^ Many proteins feature disordered regions besides folded domains, while other proteins are completely unstructured. Some intrinsically disordered proteins (IDPs) transiently sample structures and some fold upon binding partners, while others remain unfolded even in an ultrahigh-affinity complex.^2^ Disordered regions and IDPs play essential roles in cell-signaling,^3^ where their flexibility may be vital. Assembly of IDPs in biomolecular condensates formed by liquid-liquid phase separation may be a general organizing principle in cell biology.^4^ Dysregulation of liquid-liquid phase separation and aggregation of IDPs may be the pathological mechanism in many neurological diseases. The paradigm for IDPs is arguably defined by *α*-synuclein (aS).^5^ aS was suggested to also adopt α-helical conformations throughout the whole sequence in solution in its monomeric state but recent work suggests that it is better described as a disordered random coil.^6^

Resolving the structural ensembles of IDPs in experiments is challenging due to their inherent flexibility. Nuclear magnetic resonance (NMR) studies can detect residual structures or regions in IDPs having a propensity to transiently adopt well-defined structures.^7^ Chemical shifts can provide information on secondary structure elements,^8^ which can be transiently populated, or the lack thereof as for aS.^6^ NMR J-couplings are a sensitive probe of local and in particular backbone structure.^9^ Small-angle X-ray scattering (SAXS) can complement NMR experiments by reporting on the global structures of IDPs.^10^

Generating representative structural ensembles to interpret experiments remains difficult. Data-driven models on the basis of PDB statistics, so-called coil models, have been successfully used to study disordered proteins.^11–14^ Coil models provided first insights into the molecular structure of unfolded states of proteins and IDPs. Such models, including the *flexible-meccano*^14,15^ model, have been used to interpret NMR data and also solution scattering data. NMR data can be rigorously incorporated into coil models.^6,16,17^ However, modeling based on the statistics of the *ϕ* and *ψ* backbone dihedral angles does not capture correlations between different degrees of freedom, e.g., between backbone and sidechain conformations. Coil models, while capturing the overall flexibility of IDPs, are essentially static and typically do not give insight into the dynamics of IDPs.

Molecular dynamics (MD) simulations can capture these correlations and hold the promise to resolve the structure and dynamics of IDPs with atomistic resolution. MD simulations are highly complementary to experiments.^18–20^ However, two critical issues have stymied the full power of MD simulations: (1) Inaccuracies in the force fields and (2) the inherent slow dynamics of IDPs. A myriad of shallow free energy minima for an IDP mean that any imbalance in force field will be amplified,^21^ resulting in heavily skewed conformational ensembles. Due to the countless minima, simulations will relax slowly and only a fraction of the conformational space will be visited in a typical MD simulation. Force fields for IDPs have seen a lot of development.^22–24^ In particular, dispersion-corrected water models^25^ have led to better solvation and more realistic simulations of IDPs. Simulations of small disordered systems demonstrated that local structure can be captured very well in all-atom molecular dynamics simulations with explicit solvent.^26,27^ In comparison to the dramatic improvements in hardware and software, less progress has been made on overcoming the issue of slow relaxation of IDPs associated with their large conformational entropy. Their inherent disorder and the resulting difficulty in describing their structural states and relatively large size render applications of enhanced sampling methods^28^ such as umbrella sampling, metadynamics^29^ and replica exchange molecular dynamics challenging. These methods are tailored primarily to overcome energetic barriers, and less to sample the vast and weakly structured energy landscape of an IDP.

A promising avenue to overcome the sampling limitations in molecular simulations of IDPs would be to judiciously start simulations from representative starting configurations. By choosing relevant starting configurations, rather than a single starting structure, as is typically done, one can (1) obtain much better overall sampling^30^ for a given amount of computer time and (2) exploit the parallelism afforded by large-scale computing resources^31^ such as Folding@Home, cloud computing and supercomputers. Running many appropriately initialized simulations rather than a single long simulation makes better use of available computing resources, by overcoming limits in the scaling of MD engines to a large number of cores. For simulations of folded proteins, automated ways to generate simulations from all available experimental structures and homology models based on experimental structures of related sequences have been developed.^32^ Alternatively, enhanced sampling simulations have been used to generate useful starting configurations for MD simulations.^33^ In simulations of disordered polymer melts^34,35^ and biological membranes,^36^ multi-scale approaches have proved to be very successful. Coarse-grained simulations are used to explore the space of possible arrangements. Equilibrated structures from coarse-grained simulations can then be used as starting points for simulations with more accurate all-atom force fields.

An efficient way of sampling possible three-dimensional arrangements of polymer chains is provided by chain-growth Monte Carlo algorithms.^37^ In chain-growth algorithms, the polymer chain is assembled from structures of fragments of the full-length chain. The fragments, which are small by construction, can be sampled with accurate but computationally expensive methods. Many different full-length arrangements^38^ can then be sampled by using, e.g., a coarse potential energy function. In a pioneering application of fragment assembly, Stultz et al. created ensembles of tau^39^ and aS^40^ biased towards NMR and SAXS data.

Here, we adopt an hierarchical algorithm to create large ensembles of full-length IDP structures. These structures can be used as starting points for MD simulations and for ensemble refinement against experimental data.^17,27^ We first perform all-atom MD to create extensive ensembles of fragment structures. We then merge the structures of fragments overlapping along the sequence to assemble full-length structures. The number of assembly steps grows only as the logarithm of the IDP length. This logarithmic dependence and the imposition of steric exclusion at every step of the assembly ensures a high computational efficiency of the hierarchical assembly. We show that the resulting ensembles are much more diverse than ensembles sampled in MD runs at comparable computational cost. Moreover, each structure entering the ensemble provides an excellent starting point for large scale parallel all-atom simulations on high performance computing (HPC) resources. Interestingly our ensembles agree well with high-resolution information from experiment without further refinement, emphasizing that our ensembles are useful for a direct structural analysis and as starting points for MD.

## THEORY

### Self-Avoiding Random Walk

Chain-growth Monte Carlo affords to connect accurate descriptions of local structure with exhaustive sampling of global structure^37,38^ (Figure 1). Consider recursive growth of a heteropolymer chain with steric exclusion from a set of fragments. Note that interactions other than excluded volume could be considered,^38^ but for simplicity we focus on steric exclusion. Let *i_k_* be the index of the fragment structure at segment *k* of the polymer and *c_k_* the number of such fragments. Then the structure of the polymer up to step *n* is uniquely described by the sequence (*i*_1_*i*_2_ … *i_n_*) with *i_k_* ∈ {1,…, *c_k_*}. Note that for a heteropolymer the fragments will be different.

**Figure 1:**
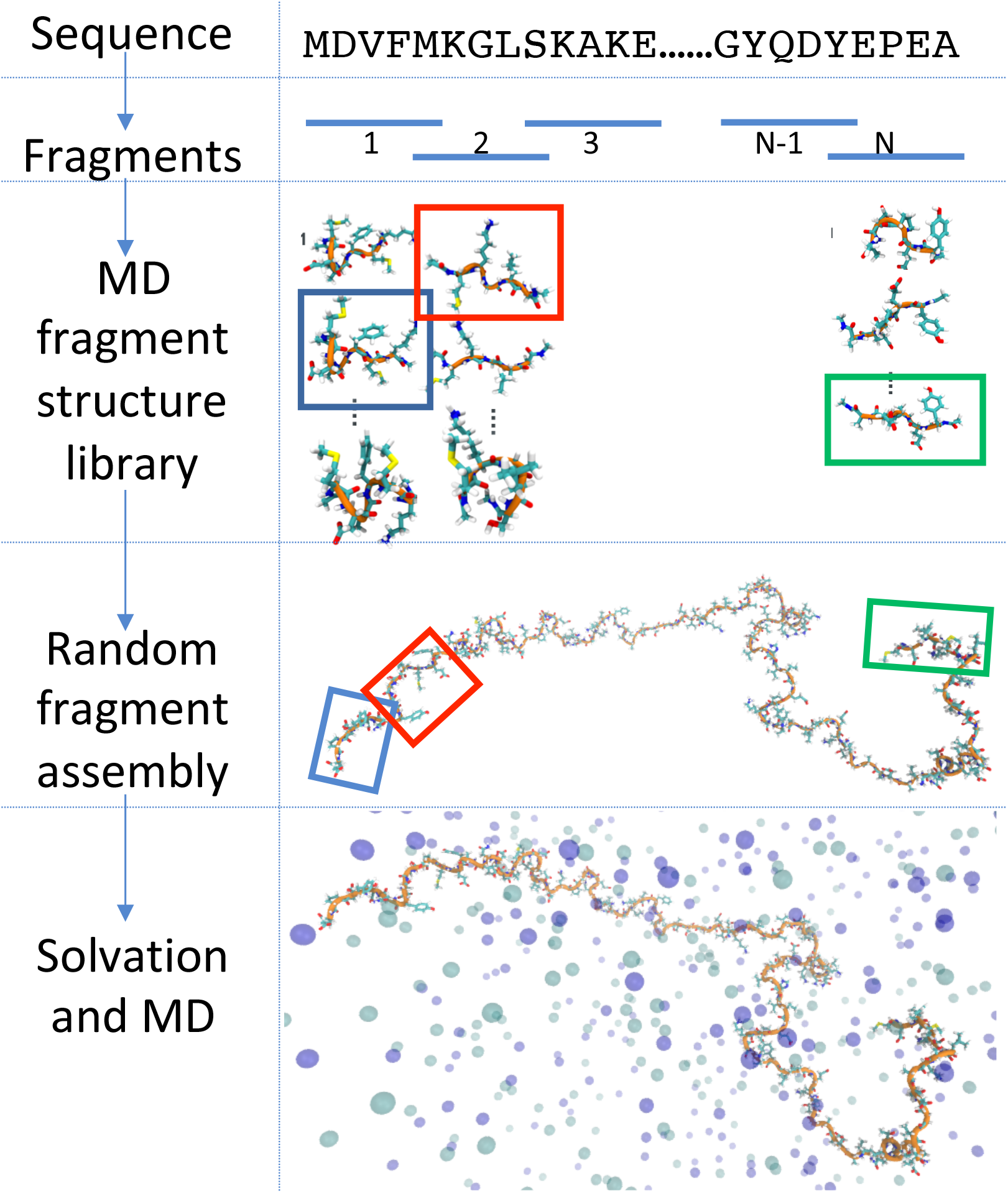
Overview of the hierarchical chain-growth approach to construct models of disordered proteins with atomic resolution in MD simulations. The sequence of the full-length protein is split into overlapping fragments, which can be sampled extensively. Fragment structures are assembled in an hierarchical manner by chain-growth Monte Carlo, sampling the space of possible global structures. The resulting full-length structures have atomic resolution and serve as starting points for highly parallel all-atom MD simulations.

The partition function of the polymer composed of *N* fragments is

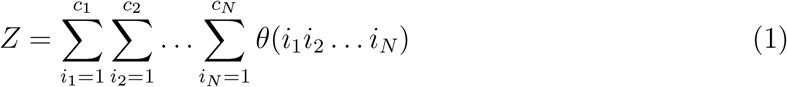

where *θ*(*i*_1_*i*_2_ … *i_N_*) = 1 if the chain is sterically permitted and *θ*(*i*_1_*i*_2_ … *i_N_*) = 0 otherwise. The probability of a particular configuration *i*_1_*i*_2_ … *i_N_* is then

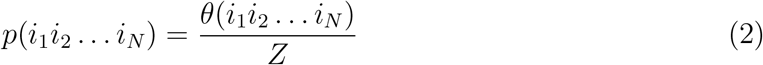

### Merging Chains from a Non-Overlapping Sub-Ensemble Using an Hierarchical Approach

For long chains we take advantage of the fact that the problem is hierarchical. We first define a segment partition function,

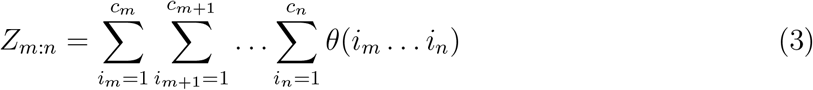

for 1 ≤ *m* < *n* ≤ *N* such that *Z*_1:*N*_ ≡ *Z_N_*. The probabilities of two sub-chains of length *k* and *N* – *k* with 1 < *k* < *N* with fragment sequences (*i*_1_*i*_2_…*i_k_*) and (*i*_*k*+1_*i*_*k*+2_…*i_N_*) can be written as

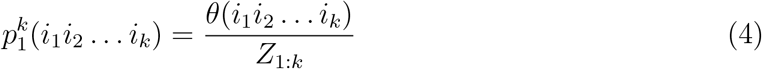

and

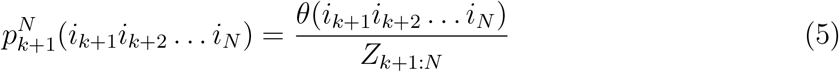

where 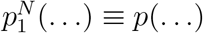 as defined above.

In the merge step, one conformation each from the ensembles of clash-free sub-chains is drawn with probabilities 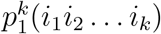 and 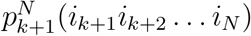, respectively. The two subchains are merged if this does not lead to a clash, i.e., if *θ*(*i*_1_*i*_2_ … *i_k_, i_k_*+_1_*i*_*k*_+_2_ … *i_N_*) = 1. The normalized probability of the merged chain then satisfies

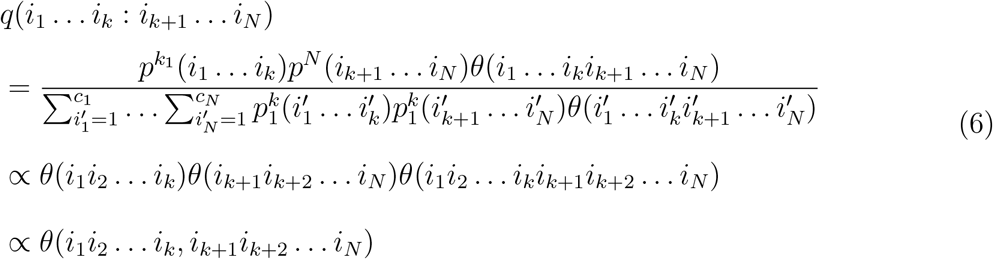

In deriving the final probability we exploited the fact that if *θ*(*i*_1_*i*_2_…*i_k_*, *θ*_*k*+1_…*i_N_*) = 1 then *θ*(*i*_1_*i*_2_…*i_k_*) = *θ*(*i_k_* + 1*i*_*k*+2_…*i_N_*) = 1. I.e., if there is no clash in the full-length chain, there cannot be a clash in any of its sub-chains. From eq 6 it follows that, properly normalized,

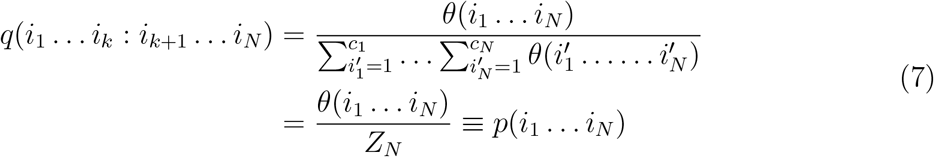

I. e., if two chains are picked at random from the 1: *k* and *k* + 1: *N* ensembles, respectively, and then merged without a clash, they enter the 1: *N* ensemble with uniform weight. In essence, merging two sub-chains without re-weighting is possible because θ ∈ {0,1}. If we instead had weight factors defined by a Boltzmann factor for a potential energy that varied continuously, instead of assuming only values *U* ∈ {0, ∞}, then we would need to reweight the merged chains.^38^

### Hierarchical Algorithm to Generate Self-Avoiding Random Walks

The ability to merge sub-chains makes it possible to grow chains hierarchically. This procedure is particularly efficient if the number of fragments *N* is a power of 2, i.e., *N* = 2^*M*^ with *M* integer. We can then obtain properly weighted chains of length 2*L* by merging chains of length *L* and checking for clashes. Starting with monomers (*L* = 1) we end up with full-length chains after *M* steps. We first create ensembles of sterically allowed pairs (*i*_1_*i*_2_), (*i*_3_*i*_4_), … (*i*_*N*–1_*i*_*N*_) (Figure 2A). Then we create ensembles of allowed quadruplets as pairs of allowed pairs, (*i*_1_… *i*_4_) = ((*i*_1_*i*_2_), (*i*_3_*i*_4_)) etc., checked for steric clashes (Figure 2B). From these ensembles, we create ensembles of sterically allowed octuplets (Figure 2C) as pairs of sterically allowed quadruplets. At the *M*-th step, we obtain the final structures with the proper weight in the ensemble of sterically allowed structures (Figure 2D). Chains can also be merged hierarchically if *N* is not a power of 2. Instead of factorizing by larger prime numbers and merging, say, triplets, here we consistently merge pairs. The merged sub-chains can then differ in length. At every step, one merges pairs where possible and otherwise promotes the remaining singlet, as sketched graphically in Figure S1. This procedure requires *M* steps where 2^*M*-1^ < *N* ≤ 2^*M*^.

**Figure 2:**
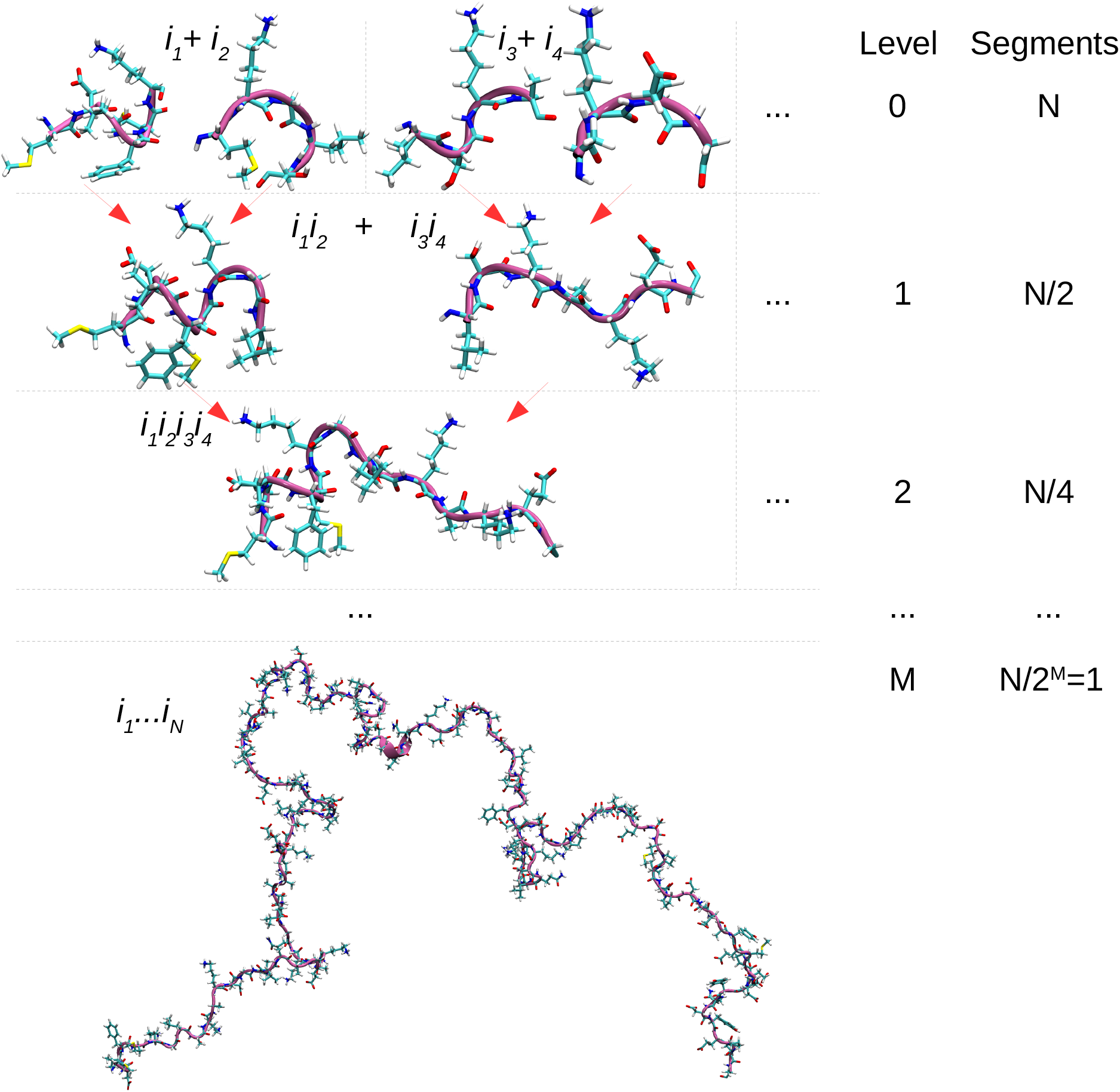
Generating all-atom IDP structures using an hierarchical Monte Carlo chain-growth algorithm. Level 0: To generate input for the chain growth, the full-length protein is split into *N* fragments, which are thoroughly sampled in all-atom MD simulations. Here, each fragment has an overlap of two residues with the subsequent fragment. Structures from the fragment libraries are picked at random to generate pairs of fragments. *N* depends on the chain length with *N* ≤ 2^*M*^, where *M* is an integer (here: *N* = 2^*M*^). Level 1: Create ensembles of *N*/2 quadruplets. Level 2: Create ensemble of *N*/4 quadruplets and so on until, at level *M*, one arrives at a full-length structure of the IDP. In each fragment-assembly step, we enforce excluded-volume interactions.

## SIMULATION METHODS

### Implementation of Hierarchical Chain-Growth Monte Carlo Algorithm

We assembled the fragments form temperature replica-exchange molecular dynamics (REMD) simulations^41^ into full-length structures, as illustrated in Figure 1, using the hierarchical chain-growth Monte Carlo algorithm (Figure 2) described above. Here, each fragment had an overlap of two residues with the subsequent fragment and capped termini (Figure 3A), but other choices are possible. Only steric interactions between fragments were considered in the chain growth. The chain-growth algorithm was implemented by building on the MDAnalysis Python library^42,43^ as described below.

0. We randomly draw two conformations from the whole set of sampled conformations in the preceding hierarchy level.
1. We perform a rigid body superposition over the peptide bonds between residues *j* – 1 and *j* of the first fragment and between residues *i* and *i* + 1 of the subsequent fragment. Here, *i* designates the first and *j* the last residue of a fragment (excluding the end-capping groups). Thus, we align the four backbone atoms C, O, N, and H in the peptide bonds between residues common to both fragments (Figure 3A, residues 4 and 5).

a. If the root-mean-square deviation (RMSD) of the superimposed region is below a given cut-off, here 0.6 Å, we accept the alignment.
b. Else we discard the conformations, draw new conformations and start again with step 1.
2. We check the aligned structures for clashes. The excluded volume is detected by calculating a neighbor list (as implemented in MDAnalysis^42,43^) for the residues from both fragments outside the alignment regions. I.e., all atoms from residues *j* — 1 and *j* of the first and residues *i* and *i* + 1 of the subsequent fragment as well as hydrogen atoms are excluded from the calculation of the neighbor list. Heavy atoms within a distance of 2.0 Å count as clash.

a. If no steric clash was detected we proceed to merging the fragments.
b. Else we discard the conformations, draw new conformations and start again with step 1.
3. We stitch the superimposed fragments together. We merge the fragments by taking backbone and sidechain atoms of residues *i*: *j* – 1 from the first fragment and *i* + 1: *j* from the subsequent fragment. In Figure 3B, the alignment, clash calculation and the stitching procedure is shown exemplary for hierarchy level 0 fragments 0 and 1.

**Figure 3:**
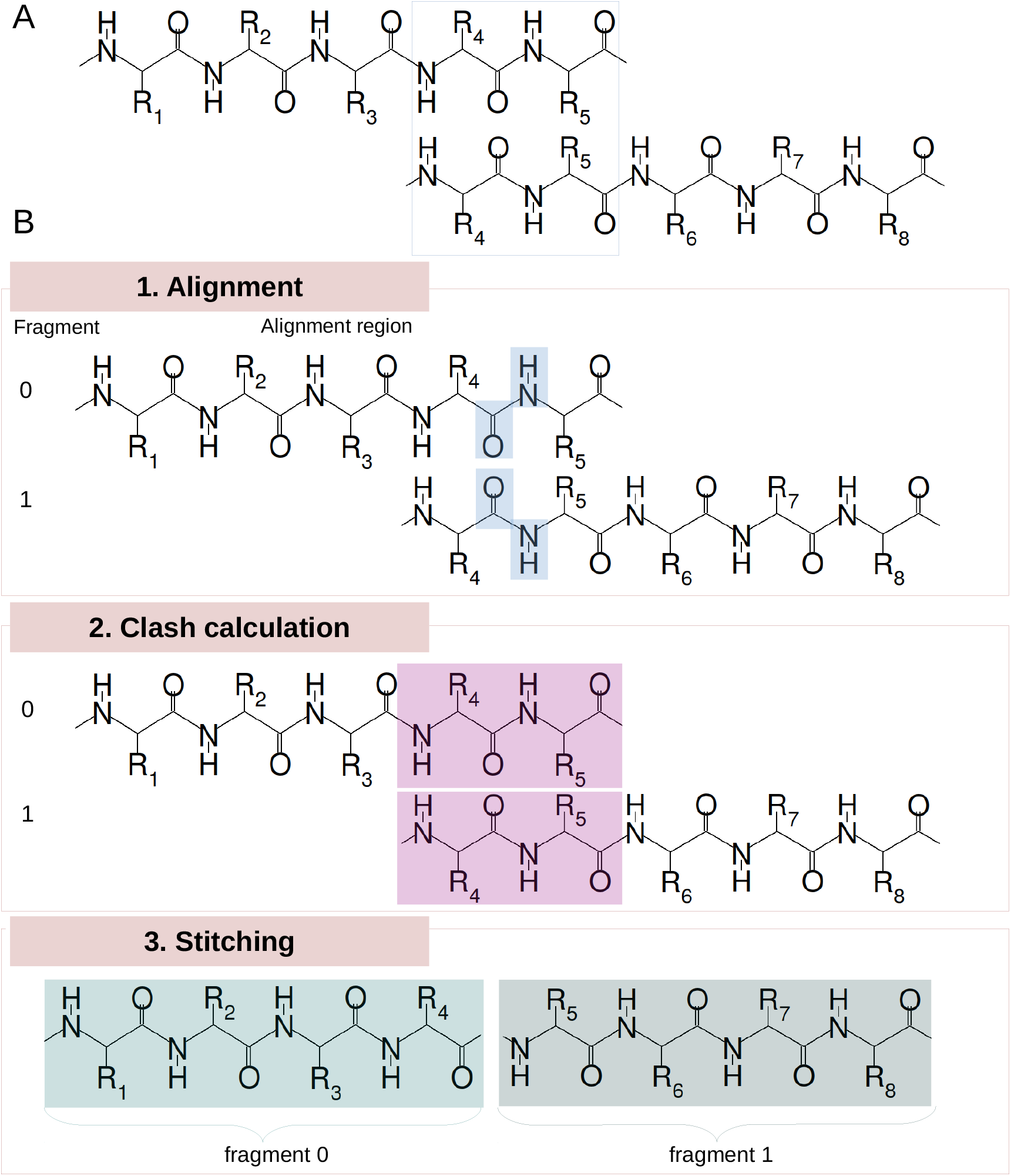
Implementation of the hierarchical Monte Carlo chain-growth algorithm. The algorithm is illustrated for the merging of two fragments at the beginning of a peptide chain (i.e., fragments 0 and 1 at hierarchy level 0). (A) Fragments with a sequence overlap of two residues with the subsequent fragment (blue box) are used as input. (B.1) To merge two randomly picked fragments at hierarchy level 0, the peptide bonds (blue shading) of the fragments are aligned. If the RMSD of the superimposed atoms is below a cut-off (here 0.6 Å), the aligned fragments are checked for clashes. (B.2) Steric overlap is probed with a heavy-atom cut-off distance of 2.0 Å. Residues (backbone and sidechain) at the alignment point (magenta shading), the endcapping groups (ACE and NME), and hydrogen atoms are excluded from the clash calculation. (B.3) If no steric clash is detected, the structure combining residues 1-4 from fragment 0 and residues 5-8 from fragment 1 is stored for use in the next hierarchy level.

### REMD Simulation of Fragments

REMD simulations were run in GROMACS/2016.4^44^ with the AMBER99SB*-ILDN-q force field^26,45–47^ and the TIP3P water model.^48^ The 46 aS penta-peptide fragments were capped at the N- and C-terminus by acetyl and N-methyl groups, respectively. The fragments were solvated in water with 150 mM NaCl, ensuring overall charge neutrality. The resulting systems contained about 1900 atoms each. For each of the 46 aS penta-peptide fragments, REMD simulations were performed for 100 ns using 24 replicas spanning a temperature range of 288 K to 431 K at constant pressure, at temperatures set according to the algorithm by Patriksson *et al*.^49^

To maintain a pressure of 1 bar, the Parrinello-Rahman^50^ barostat was used. Temperature coupling was achieved by velocity rescaling with a time constant of 0.1 ps using the Bussi-Donadio-Parrinello thermostat.^51^ The P-LINCS algorithm was used to constrain all bonds.^52^ Using the particle mesh Ewald method, long-range electrostatics were calculated with a cut-off of 10 Å. The van der Waals cut-off was set to 12 Å. REMD production runs were preceded by energy minimization and 1 ns equilibration in the NPT ensemble. During the production runs of 100 ns (per replica) structures were saved in intervals of 10 ps. In this way, we created a library of 10 000 fragment structures for each peptide segment at the temperature of interest, *T* = 288 K.

### MD Simulations of Full-Length Models

All-atom MD simulations of full-length aS were run in Gromacs/20 1 6.4^44^ with the AMBER99SB*-ILDN-q force field^26,45–47^ using the TIP4P-D water model.^25^ Twenty models were chosen at random from the ensemble of models generated by hierarchical assembly. The aS chains with charged termini were each solvated in water with 150 mM NaCl, ensuring overall charge neutrality. Each system contained about 350 000 atoms. Simulations were performed at a constant temperature of 300 K using the Bussi-Donadio-Parrinello velocity-rescaling thermostat with a time constant of 0.1 ps.^51^ The pressure was maintained at 1 bar using the Parrinello-Rahman barostat.^50^ The P-LINCS algorithm^52^ was used to constrain all bonds. To calculate long-range electrostatics the particle mesh Ewald method was used with a cut-off of 12Å. A cut-off of 12 Å was used for van-der-Waals interactions. Energy minimization and 200 ps equilibration were performed before running production runs of 100 ns.

### Calculation of Experimental Observables

We calculated NMR chemical shifts with SPARTA+.^53^ Reference random coil chemical shifts were predicted using the POTENCI web server developed by Nielsen and Mulder^54^ to arrive at secondary chemical shift predictions by subtraction of the reference value. J-couplings were calculated as previously described^55^ using the original Karplus parameters described by Wirmer and Schwalbe.^56^ The radius of gyration *R_G,i_* for a saved aS structure *i* was calculated using FoXS,^57^ taking the solvent shell into account. In addition, a geometric radius of gyration *R_G,i_* was computed from the protein coordinates with the MDAnalysis Python library.^42,43^ The model ensemble average was calculated as the root-mean-square average, 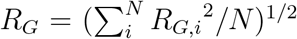, where *R_G,i_* is the radius of gyration of the *i*-th member of the ensemble of size *N*.

## RESULTS AND DISCUSSION

### Testing and Validation of the Hierarchical Chain-Growth Approach

We first verified that the Monte Carlo hierarchical chain-growth algorithm works as expected and tested different practical implementations. In the hierarchical chain-growth algorithm, we assemble the full-length chain by generating possible structures of dimers of fragments (Figure 2A) and then the structures of possible quadruplets (Figure 2B) and so on until the full chain is grown (Figure 2D). As expected from its derivation, the chain-growth algorithm generates ensembles in which each structure appears with a Boltzmann weight, which we confirmed by comparing chains with up to 26 residues to chains grown by brute-force generation of self-avoiding random walks (Figure S2). We also evaluated the effect of using fragments of different length to grow 50-amino-acid long aS sub-chains. Comparing 3mer, 4mer and 5mer fragments, we found that using larger fragments resulted in somewhat more compact models (Figure S4A), with a 5mer having a radius of gyration *R_G_* about 1 Å less than a 4mer. Larger fragments can adopt more compact conformations, with more interactions within the fragments. In terms of the end-to-end distance χ (Figure S4B) and the diversity of structures, as measured by the pairwise RMSD between structures in the ensemble (Figure S4C), the differences between 3mer, 4mer and 5mer fragments are small. We decided to use 5mer fragments in the following to capture also less-extended structures in our initial pool of fragment structures.

Secondly, we compared the effects o aligning the fragments at the peptide bond (Figure 3) or the backbone of the residue at the merge point (Figure S3). The peptide bond is relatively rigid due its partial double-bond character compared to the backbone of the residue at the merge point with its rotatable *ϕ* and *ψ* dihedral angles. Indeed, using the backbone rather than the peptide bond for alignment results in larger differences between the assembly of 5mer, 4mer and 3mer fragments, as judged by the distributions of *R_G_, χ* and pairwise RMSDs between structures (Figure S4). Importantly, assembling the chain via alignment of the peptide bond preserves the conformational distributions from MD simulations of the fragments. Figure 4A and B illustrate this point for the *ψ* angle of A11 and Y39. By contrast, assembly via the backbone introduces a bias towards extended structures with *ψ* > 100°.

**Figure 4:**
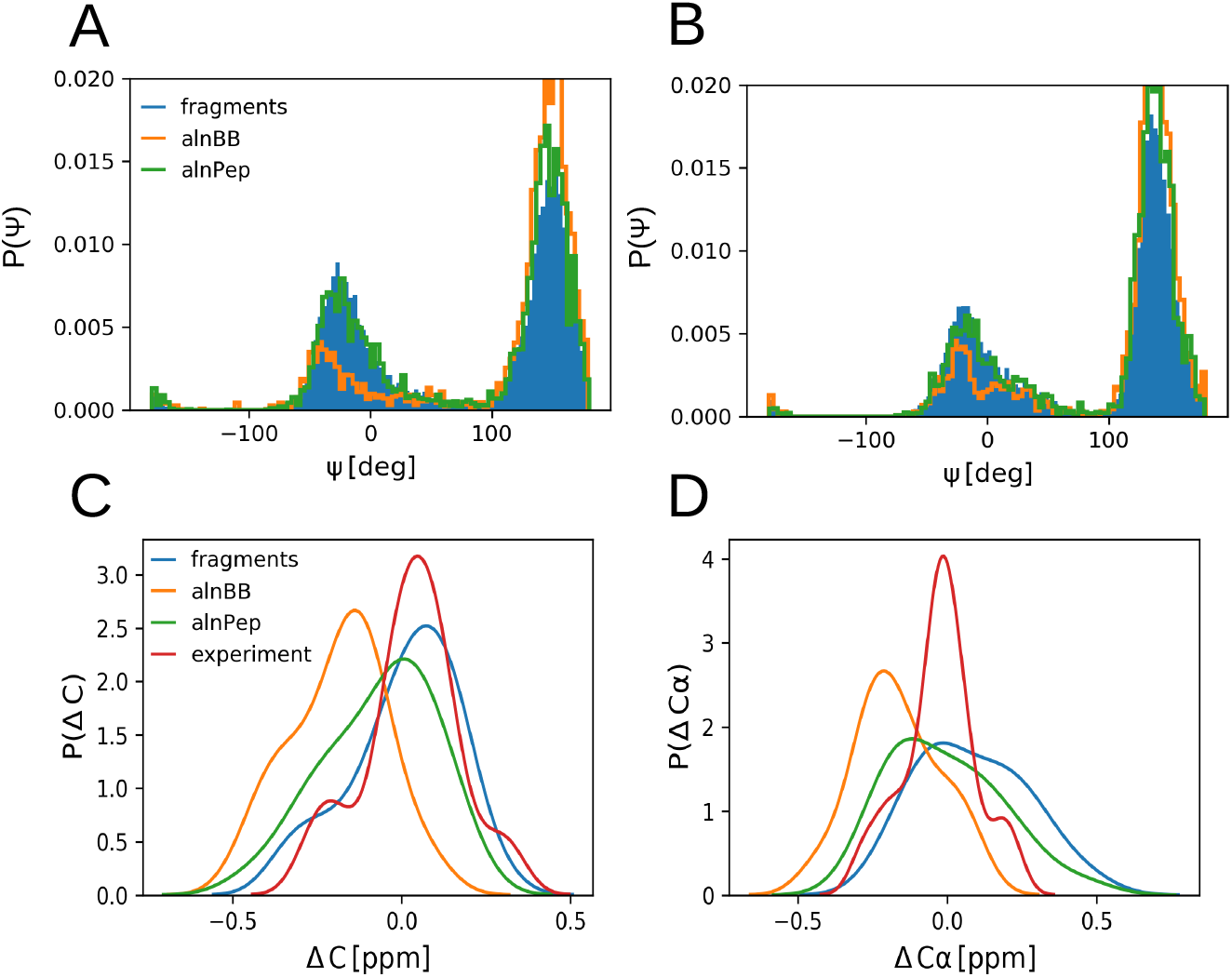
Comparison of different alignment and stitching approaches to implement the chain-growth algorithm. (A,B) Distribution of the *ψ* dihedral angle for A11 (A) and Y39 (B) before and after the assembly of the MD fragments into full-length structures. Results are shown for two different alignment approaches (orange: alignment over the backbone atoms, alnBB; green: alignment over the peptide bond, alnPep) and for the peptide fragments (blue). (C,D) Distribution of Δppm secondary chemical shifts of ^13^C (C) and ^13^Cα (D) from experiment (red), MD fragments (blue), and full-length models grown with backbone alignment (orange, alnBB) and peptide bond alignment (green, alnPep).

Thirdly, we considered different overlap between the fragments. Using an overlap of one or two residues between the fragment makes no significant difference (Figure S5). For subsequent chain growth, we used the central three residues of 5mer fragments. The central residues of a fragment should be more representative of the local structure in the context of an IDP chain.

We conclude that the different choices one could make in implementing our chain-growth algorithm, overall, do not have drastic effects on the global structures of the generated ensembles. Local structure may be preserved better by aligning on the rigid peptide bond rather than on the more flexible backbone.

### Full-length Models of aS

We generated a highly diverse ensemble of full-length allatom aS structures with the hierarchical Monte Carlo chain-growth algorithm. For aS, we split its primary sequence into 46 fragments to produce input structures for the chain growth. For each fragment we ran exhaustive atomistic simulations with explicit solvent using REMD. This initial sampling phase already lends itself to HPC resources with a large number of computing nodes as each fragment can be simulated independently from the other fragments and no overhead is incurred due to inter-node communication. During chain growth, structures are drawn from the simulation ensembles for the individual fragments.

Using the hierarchical chain-growth algorithm described above we grew 20 000 full-length aS models. The calculation ran on 20 compute cores and we grew 1 000 full-length models per core using the same pool of fragments in the 20 runs but different random number seeds. To sample an ensemble with a large diversity in local conformations in the highest level *M*, we grew 10 000 structures in the levels 0 to *M* — 1. The extensive sampling of local and global structures yielded highly-diverse full-length structures of aS, as judged by pairwise RMSD (Figure 5D), with an average pairwise RMSD of ≈56 Å between the 20 000 models.

**Figure 5:**
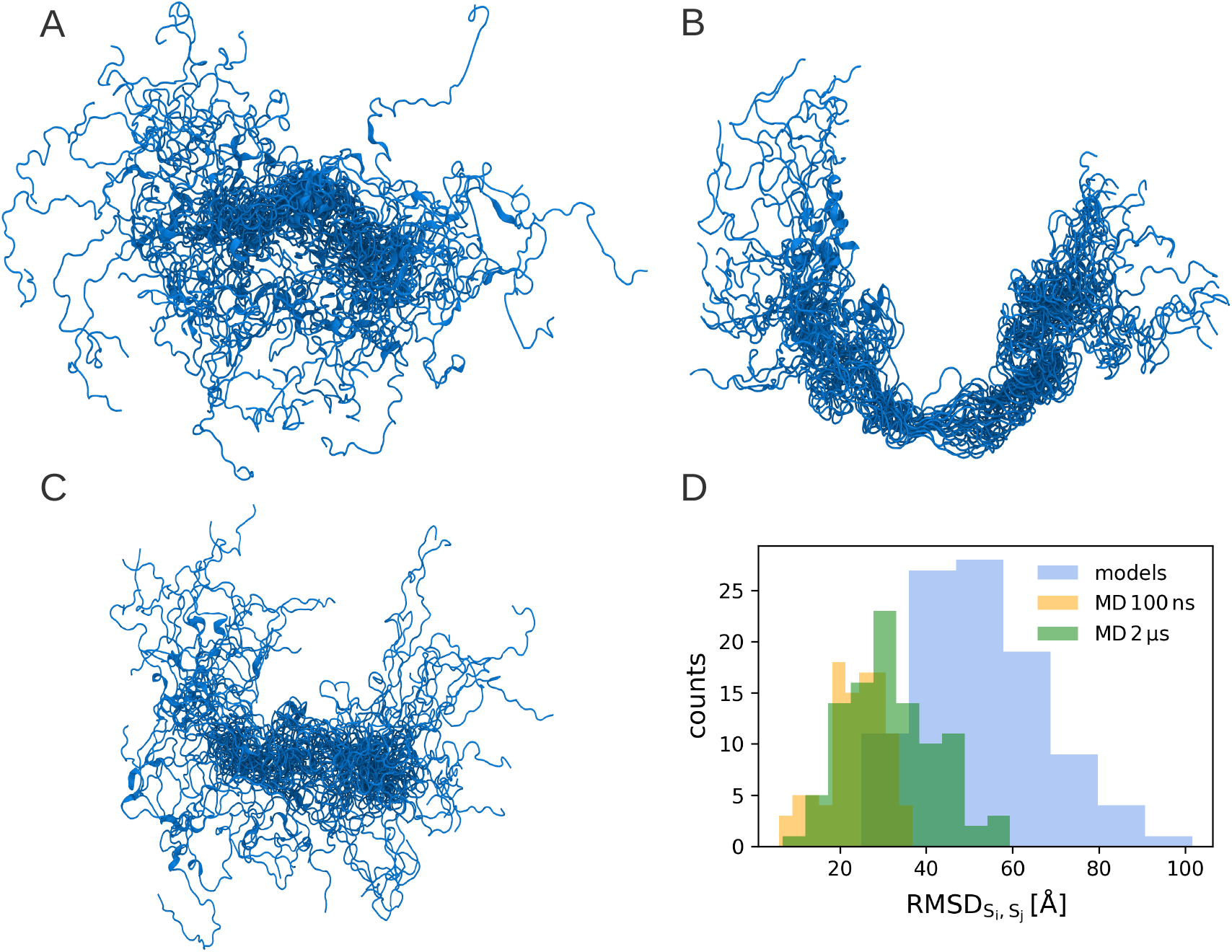
Conformational diversity of aS model ensembles from MD simulation and hierarchical chain growth. (A) 30 aS models from hierarchical chain growth. (B) 30 structures from 10 ns of MD. (C) 30 structures from 100 ns of MD. The structures were aligned on the central third of their sequence in (A), (B) and (C). (D) Distribution of pairwise RMSD between 100 different models obtained by MD, sampled uniformly in time from 100 ns and 2 μs, respectively, and by hierarchical chain growth.

In essence, no two structures are alike.

### Sampling Efficiency of the Hierarchical Approach

Visualization makes it clear that our hierarchical approach captured a much larger conformational space than would be accessed in a typical all-atom MD simulation of an IDP. In Figure 5B the persistence of a transient hairpin conformation over the course of 10 ns of MD simulation is clearly visible. Such local structure decorrelated over 100 ns of simulation (Figure 5C). However, it is clear from comparing Figure 5A and Figure 5C that the ensemble from chain growth sampled a much larger conformational space. The structures from 100 ns of MD simulations still re-sembled one another, unlike the structures from chain-growth, which fully explore the space of possible structures.

We compared the distribution of RMSD values between 100 models sampled with hierarchical chain growth to 100 different conformations sampled in a 100 ns and a 2 *μ*s MD simulation. The 100 structures from the 100 ns and 2 *μ*s MD simulation trajectories were taken at regular time intervals of 1 ns and 20 ns. For the chain growth ensemble with 100 different models we observed an average pairwise RMSD of ≈53 Å, whereas the MD ensemble after 100 ns showed an average RMSD of ≈23 Å and after 2 *μ*s an average of ≈32 A (Figure 5D). This demonstrates that by using the hierarchical chain-growth algorithm we were able to obtain a diverse ensemble, much more diverse than an ensemble sampled in 2-*μ*s of MD.

Principal component analysis (PCA)^58^ gives us a global view of the conformational space sampled. Figure 6 shows projections of the 20 000 models constructed with hierarchical chain growth onto the principal components 1 and 2. On top, Figure 6A shows 200 000 MD conformations sampled in 20 × 100 ns of MD simulation started from 20 different structures (black crosses). In essence, each of the 20 runs explores a small region within the confines of the space sampled by the hierarchical models. Figure 6B shows the 2 *μ*s MD simulation projected onto the principal components 1 and 2 of the hierarchical models. Again, the MD ensemble is contained within the hierarchical ensemble, indicating the larger conformational diversity of structures in the hierarchical ensemble.

**Figure 6:**
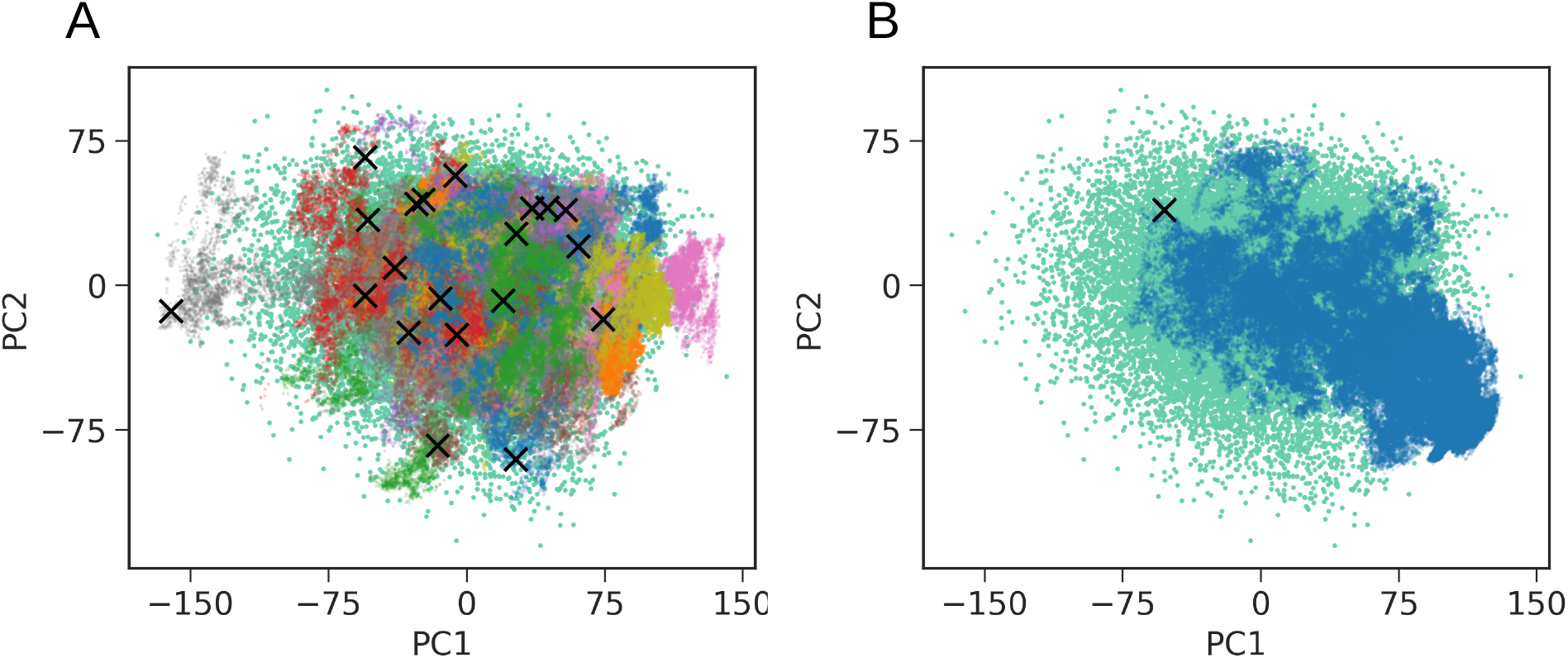
Conformational space of full-length models of aS projected onto the principal component axes 1 and 2, as obtained from the ensemble of hierarchical models. (A) Conformations sampled in 20 × 100 ns MD simulations with different starting structures (black crosses) projected onto the conformational space spanned by 10 000 full-length aS models (aquamarine). (B) 2 *μ*s trajectory (blue) projected onto the conformational space sampled by the aS models (aquamarine).

Overall, the MD trajectories stayed within the boundaries of the conformational space defined by the ensemble from chain growth, which suggests that the hierarchical sampling covered, at least at the global level of principal axes 1 and 2, the conformation space visited in the 20 × 100 ns and *μ*s scale MD simulations. Thus we conclude tentatively that our hierarchical approach exhaustively samples the global structures of IDPs such as aS.

### All-atom MD Simulations of aS in Explicit Solvent

Our models are meaningful starting points for simulation, as shown by the overall behavior of the simulations started from our models and the conservation of their characteristics in MD simulations. As expected, the full-length structures of aS (Figure 2F) were dynamic in all-atom MD simulations in explicit solvent. In Figure 7B the starting structure for an all-atom simulation in explicit solvent is presented. The structure is extended and this particular structure features a turn close to the end of the N-terminal domain and at the beginning of the C-terminal domain of aS. Like most of our models the structure shows little in the way of well-defined secondary structure elements, featuring only a short helical segment. After 50 ns of simulation the turn at the end of the N-terminal vanished and the short helix deformed (Figure 7C). Visually, the structure expanded further. Over the next 50 ns the molecule became somewhat less expanded (Figure 7D). Preservation of their characteristics in the MD simulations suggested that the models provide meaningful starting points for simulations with the state-of-the art AMBER99SB*-ILDN-q force field^26,45–47^ and TIP4P-D water model.^25^ Interestingly, the simulations showed a slight compaction of the models, as judged by *R_G_* (Figure 7A). The mean square *R_G_* dropped from ≈40 Å (which is close to experiment, as discussed below) to ≈ 33 A during the 100 ns of MD. This compaction could be an indicator of a lack of residual structure in the models or of a poor force field and solvent model, which underestimate the solvation of the protein. However, the simulations did not access fully collapsed structures, with *R_G_* ≈ 15Å similar to a folded protein of this length, suggesting that the AMBER99SB*-ILDN-q force field and TIP4P-D water model^25^ describe IDPs well enough.^24^

**Figure 7:**
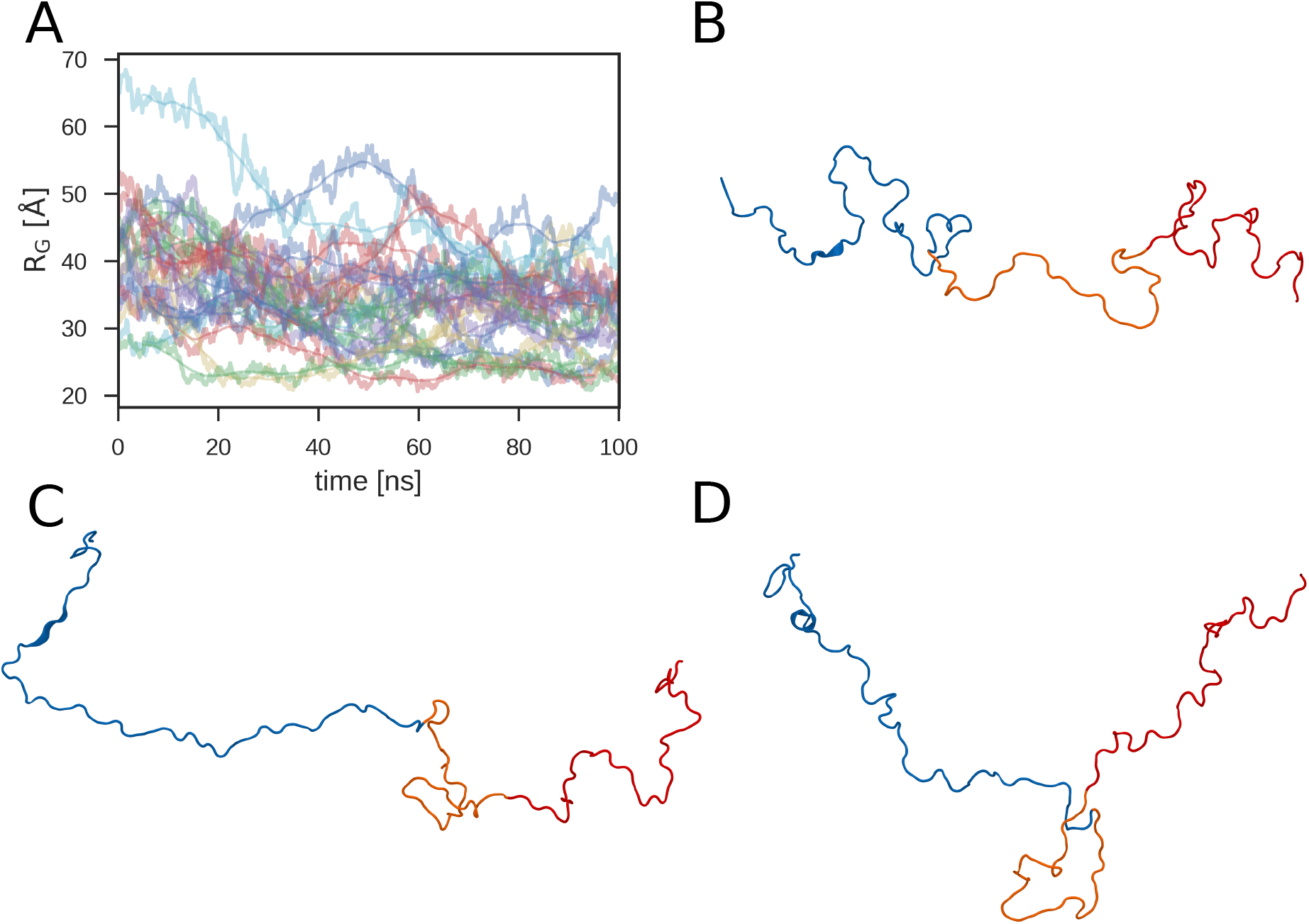
All-atom MD simulations of full-length models of aS. (A) fluctuations of the geometric *R_G_* in all-atom MD (noisy curves) and the respective moving average using a window size of 10 ns (smooth curves of corresponding color). Representative snapshots are shown for a simulation run: starting structure (B), after 50 ns (C) and 100 ns of MD (D). In (B), (C) and (D) the N-terminal domain of aS is colored in blue (M1-K60), the central region with a hydrophobic motif in orange (E61-V95), and the C-terminal domain in red (K96-A140).^59^

### Comparison of aS Ensembles to NMR Experiments probing Local Structure

The aS ensembles obtained by hierarchical chain growth compare well to NMR experiments probing the local structure through J-couplings^6^ and chemical shifts.^6,16,60^ Without reweighting, the^3^ J_CC_ and ^3^J_CHα_ couplings probing *ψ* and *ϕ* dihedral angles agree well with experiment (Figure 8A). The magnitude of ^1^J_NCα_,^2^J_CαN_ and^3^J_HNHα_ couplings, which report on the *ϕ* dihedral angles, were captured by our ensemble but small systematic offsets were observed. Values for ^1^J_NCα_ and ^2^J_CαN_ tend to be somewhat lower in our ensemble than in experiment, whereas^3^J_HNHα_ values tend to be slightly overestimated. The amide nuclear spin involved in these couplings is affected by many processes such as hydrogen-bonding or geometric distortion. These processes are not well described by current force fields, which may explain the small systematic deviations.^61^ The agreement with experiment is equally good for J-couplings calculated from full-length models (Figure 8B) and from the initial fragment library generated of short aS fragments (Figure 8A), highlighting that the chain-growth algorithm preserves the local structure sampled in REMD.

**Figure 8:**
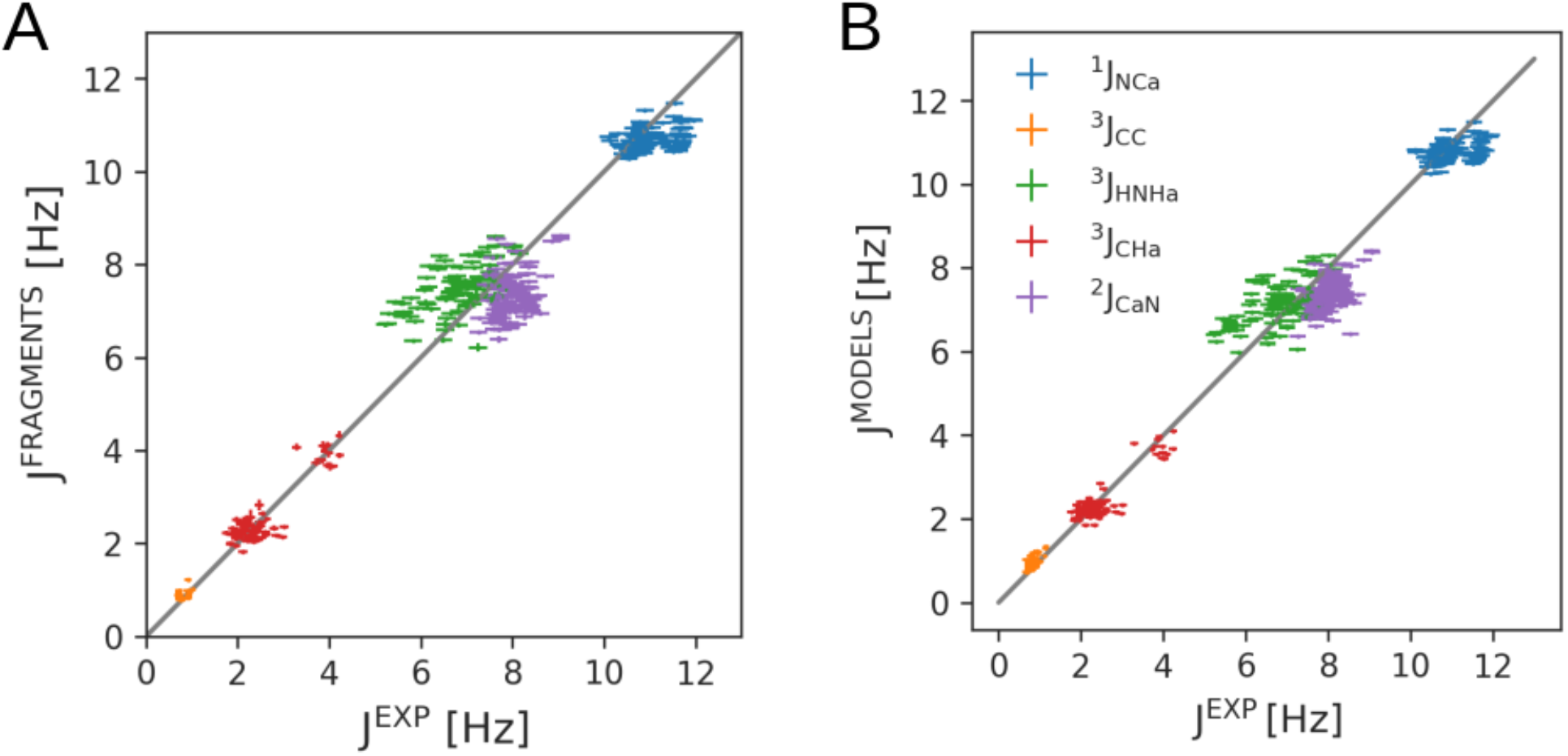
Comparison of calculated NMR J-couplings (*y*-axis) to experiment^6^ (*x*-axis). J-couplings were obtained (A) from MD simulations of peptide fragments and (B) for full-length aS chains built by hierarchical chain growth. Each point corresponds to the ensemble average of a single residue. Standard errors of the mean were estimated by block averaging.

The ^13^C and ^13^*C_α_* secondary shifts calculated from models were around zero for all residues, suggesting a local backbone conformation closely resembling random coil, in agreement with experiment^6,16,60^ (Figure 9). Deviations from zero in the secondary chemical shifts (Δppm) report on (residual) structure, but these deviations were quite small (< 1 ppm) for models compared to estimates of the expected errors in calculating chemical shifts. For empirical chemical shift prediction of ^13^C and ^13^C_α_ shifts, RMSD to experiment of about 1 ppm were found for a validation set of 11 proteins.^53^ Here, the RMSD to experiment is 0.32 and 0. 37 ppm for ^13^C and ^13^C_α_ shifts, respectively, as predicted for the fragments, and 0.33 and 0. 35 ppm as predicted for the full-length models. Indeed, for most residues the chemical shift predicted after assembly agrees with these predicted for the MD fragments before assembly. Still for few residues the experimental secondary shifts were captured better by the MD fragments before assembly to full-length models, e.g., the carbonyl secondary shifts reported for A18, Figure 9A or the Cα secondary shifts reported for V16, Figure 9B. Nevertheless for some residues the agreement with experiment was better after the assembly (carbonyl secondary shifts reported for E20, Figure 9A or the Cα secondary shifts reported for V74, Figure 9B).

**Figure 9:**
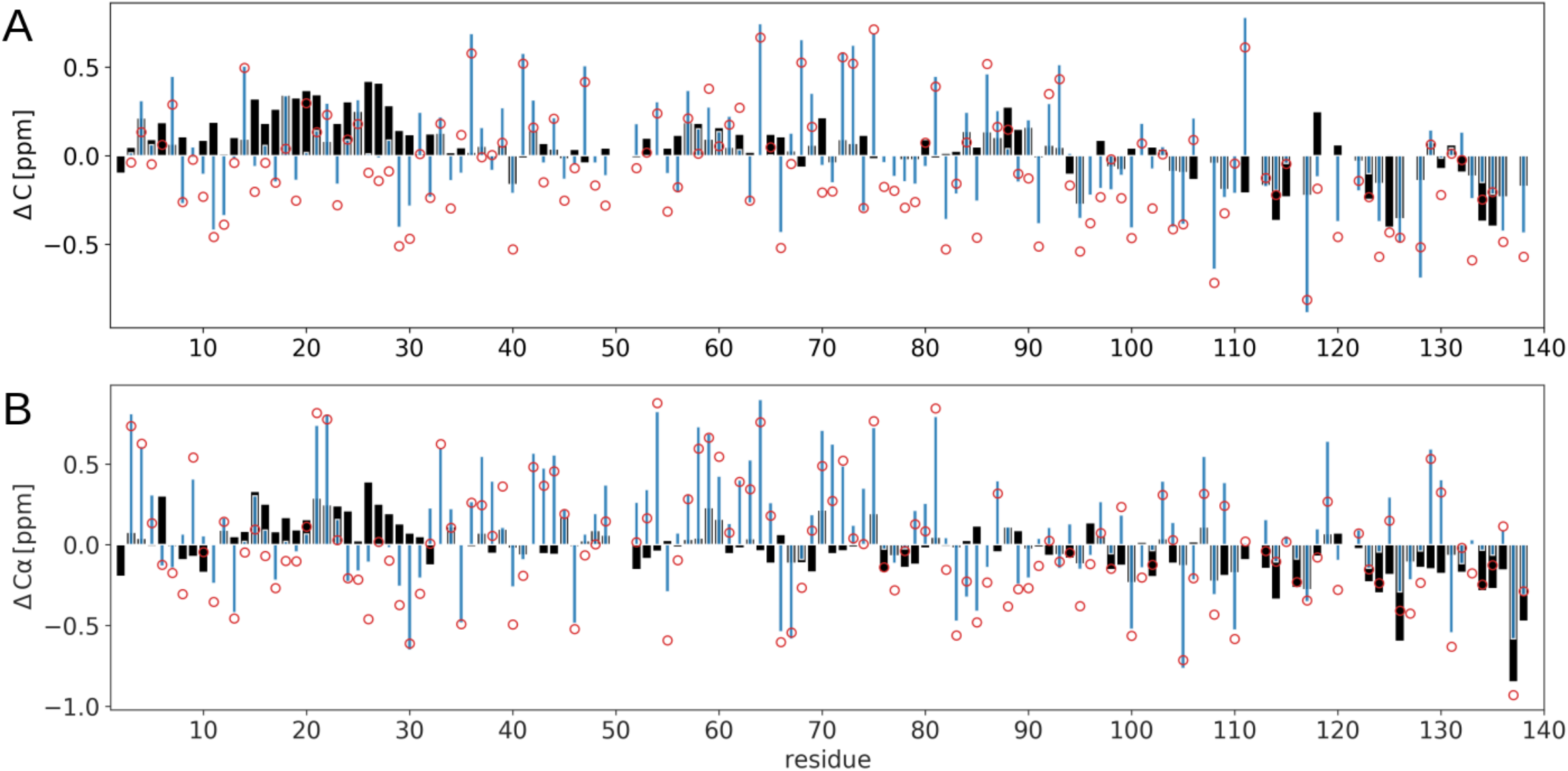
NMR chemical shift analysis for fragment simulations. (A) ^13^C and (B) ^13^C_α_ secondary shift values from fragment all-atom MD simulations (blue bars), assembled full-length models (red circles) and experiments^6,16,60^ (black bars).

In Figure 4C and D, we compared the distribution of the experimental Δppm secondary chemical shifts for the MD fragments and for full-length chains grown with different alignment procedures (alignment of the flexible backbone versus alignment of the rigid peptide bond). Growing IDP chains by aligning the backbone of the fragments at the merge point results in a shift in the chemical shift distributions, which mirrors the shift towards β-strand like conformations we found with this growth procedure (Figure 4A and B). By contrast, alignment via the peptide bond results in a much smaller shift away from the distributions from the fragment predictions (Figure S6) and experiment. Alignment via the peptide bond largely preserves the structures of the fragments as judged by chemical shift predictions, but not exactly. Naturally, full-length structures will be at least subtly different from fragment structures, e.g., some fragment structures will be sterically impossible. Taken together, the analysis of the chemical shifts demonstrates that our implementation of the chain-growth procedure leads to good models of IDP structure and conversely, that chemical shifts are useful indicators in the modeling of IDPs.

### Comparison of aS Ensembles to SAXS Experiments probing Global Structure

The global structure of the hierarchically grown aS models is also consistent with experiment, as probed by SAXS measurements of the radius of gyration (*R_G_*). The estimated root-mean-square radius of gyration for the ensemble of full-length aS models is *R_G_* 40Å (Figure 10). In calculating the *R_G,i_* values of individual structures *i*, we took the solvent shell into account using FoXS,^57^ but we obtained essentially the same *R_G_* by calculating *R_G,i_* of individual structures *i* directly from the protein coordinates. In SAXS experiments, *R_G_* values of 40Å (Binolfi et al.^62^) and 45 Å (Curtain et al.^63^) have been reported, bracketing our value. The value expected for a 140 amino acid random coil is 45 Å, using the parameters determined by Sosnick et al. considering nearest neighbor effects.^64^ We find it encouraging that the chain growth captures the overall dimensions of the disordered chain. Spurious compaction due to force field issues would require the imposition of a bias during chain growth or simulation to steer the ensemble away from overly compact structures that underestimate *R_G_*,^39,40^ or a reweighting of the ensemble.^17,27^

**Figure 10:**
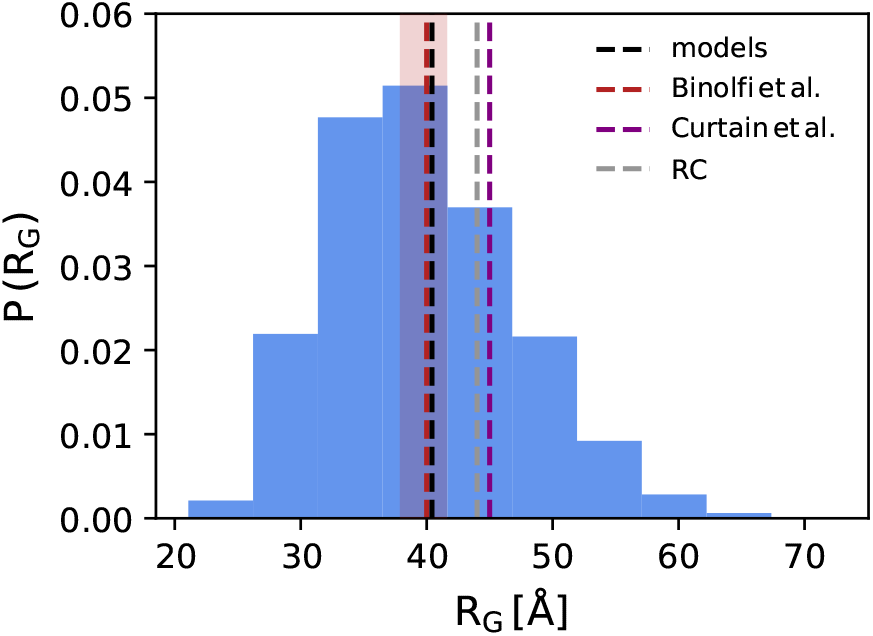
Distribution of the radius of gyration *R_G,i_* in the ensemble of aS structures obtained from hierarchical chain growth. Vertical lines indicate the root-mean-square *R_G_* value of the ensemble (black; models), *R_G_* values from SAXS experiments by Binolfi et al.^62^ (red) and Curtain et al.^63^ (magenta), and an *R_G_* estimate for a random coil (RC; gray).^64^

The observed compaction during MD, with the geometric *R_G_* dropping from 40 to 33Å in 100 ns, suggests that the chain “as grown” before MD may be a better representation of the extended structures seen in SAXS (Figure 10), with measured *R_G_* values between 40 and 45 Å. It is not unexpected that the long chains predicted to be extended undergo some kind of collapse. The factors driving the collapse, e.g., the hydrophobic effect and the potential overstabilization of protein-protein interactions, are not prominent for the very short fragments we simulated. In turn, this may indicate that the predicted chains are probably closer to the real ensemble than the MD refined ones, and that this type of setup gives us a handle to test and optimize balanced force fields for IDPs. In any case, the simulations demonstrated that AMBER99SB*-ILDN-q force field^26,45–47^ and TIP4P-D water model^25^ describe IDPs well enough, but we note that other IDP force fields may work equally well or better.^24^

Overall our hierarchically grown IDP models captured both local conformations, as reported by NMR J-couplings and chemical shifts, and the overall dimension reported by SAXS experiments, remarkably well, without any refinement. This result encourages the use and further development of our approach for generating starting conformations of highly-parallel MD simulations of IDPs and the testing of MD force fields.

## CONCLUDING REMARKS

Our hierarchical IDP chain-growth approach is perfectly suited to the exascale high-performance computing resources which are becoming available. We generated highly diverse structures, much more diverse than what one would sample in a typical MD simulation. Our structures capture both NMR data probing local structure and and SAXS data probing global structure, without any refinement, emphasizing again that the structures should be excellent starting points for MD simulations. By generating starting configurations that closely follow the Boltzmann distribution, we can launch a large number of independent simulations and this swarm of simulations can fully explore the conformational space of an IDP. Simulations setup in this way may help identify and rectify force field issues for IDPs.

It would be computationally feasible to create exhaustive fragment libraries for, say, the 20^3^ = 8000 distinct amino-acid trimers with generic flanking residues. Considering the speed of assembly, a web service for generic IDP assembly is thus feasible. It would also be possible to include post-translational modifications such as phosphorylation.

Deviations from experiment can be taken into account in a Bayesian framework. Ensemble refinement by reweighting^17^ can be applied very efficiently to large ensembles.^2^’ Bayesian analysis of all-atom simulations of a disordered peptide^27^ showed that quantitative agreement with high-resolution NMR experiments can be achieved and led to system-specific correction, which may also be important when modeling large IDPs. Stultz et al. have already shown how structural ensembles from fragment assembly can be refined against NMR and SAXS data within a Bayesian framework.^39,40^

Our approach can be extended to simulations of other flexible biomolecules and their assemblies. For instance, it could be used to model long non-coding RNAs and other singlestranded nucleic acids. For single-stranded nucleic acids, sampling is a bottleneck in force field evaluations,^65^ which could be addressed by an extension of our approach. We envisage extensions of our approach to simulate dense solutions of IDPs as in biomolecular condensates formed via liquid-liquid phase separation.^4^ Our chain-growth algorithm is valid whether individual chains or assemblies of chains are modeled. For modeling dense biomolecular condensates variants of the chain-growth Monte Carlo algorithm we employed here may prove advantageous. For more dilute condensates the current approach should lead to reasonable starting conformations for large-scale MD simulations.

## SUPPORTING INFORMATION

Implementation of hierarchical chain-growth algorithm and comparison of chain-growth algorithms, consistency checks for hierarchical chain-growth algorithm and additional analysis of predicted chemical shifts for fragments and full-length models of aS.

## Acknowledgement

We acknowledge financial support from the German Research Foundation (CRC902: Molecular Principles of RNA Based Regulation) and by the Max Planck Society. We thank Profs. Markus Zweckstetter and Harald Schwalbe and Drs. Adriaan Bax and Jürgen Köfinger for insightful discussions.

## Supplementary Text

### Self-Avoiding Random Walk

#### Implementation of Hierarchical Chain Growth for *N* not a Power of Two

As described in the main text, the hierarchical chain-growth algorithm is also applicable if *N*, the number of fragments composing the chain, is not a power of two. In Figure S1, the application of the hierarchical chain-growth algorithm to *α*-synuclein (aS) is shown, where 46 MD fragments are used to grow the full-length 140 amino-acid protein.

**Figure S1:**
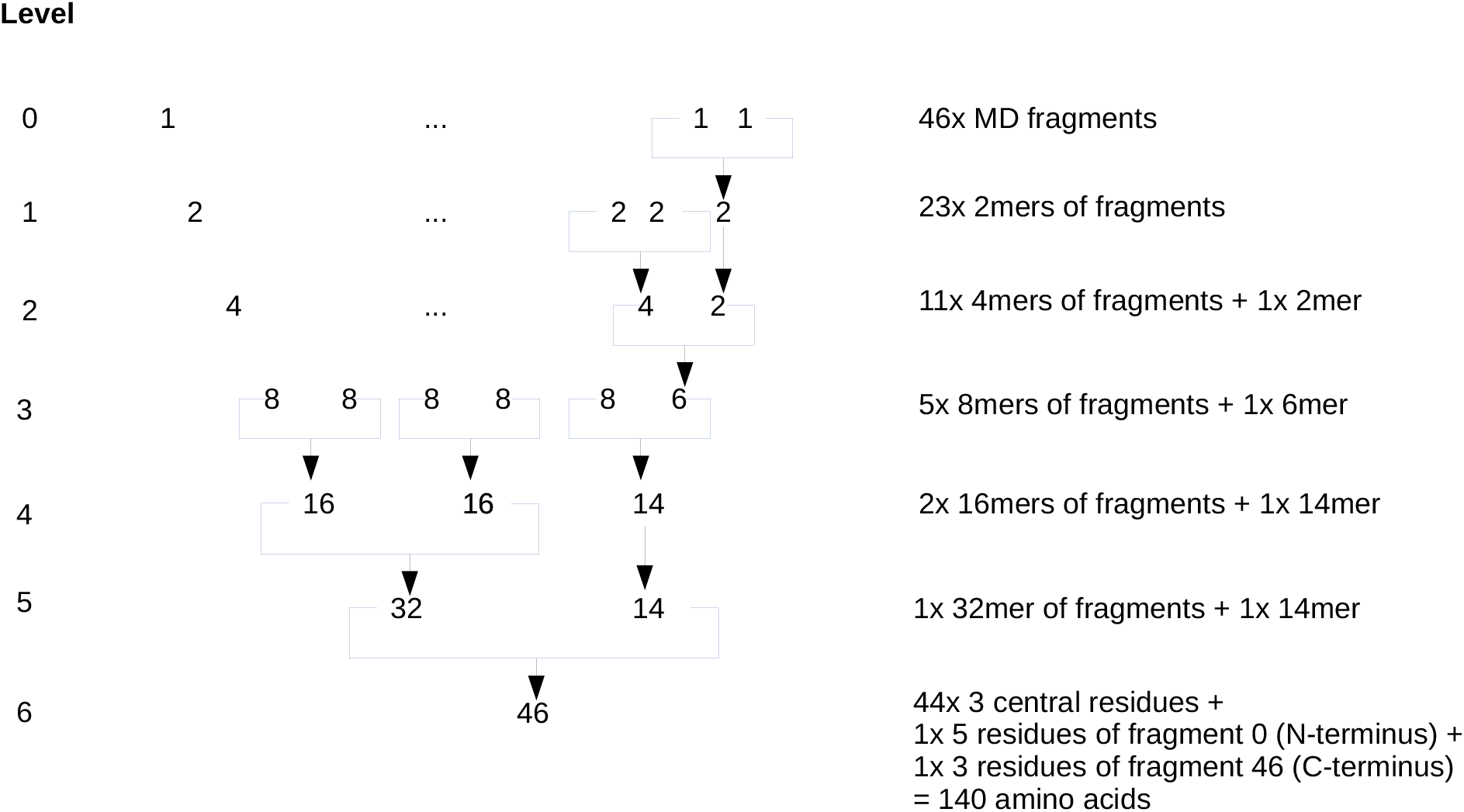
Implementation of hierarchical chain growth for aS. Here *N*, the number of fragments, is not a power of two, and the C-terminal fragment is shorter (3mer versus 5mers). At each hierarchy level, one merges fragment pairs where possible and otherwise promotes the remaining singlet to the next hierarchy level.

#### Comparison of Chain-Growth Algorithms

For reference, we also implemented a “naïve algorithm” to grow chains from fragments. In this algorithm, we draw enumerations at random and reject them if there is a clash involving any of its fragments:

1. Randomly pick a first element *i*_1_. Set *n* =1.
2. If *n* < *N*, randomly pick a new element *i*_*n*+1_; otherwise enter *i*_1_… *i_N_* into the ensemble and return to step 1 (until the ensemble has reached a certain size).
3. Check for a clash of the new element with the rest of the chain.
  a. If *i*_*n*+1_ does not clash with *i*_1_… *i_N_*, then accept the addition and increase n by one, *n* ↦ *n* +1, and go to step 2.
  b. Otherwise, go to step 1 and restart.

For long chains this algorithm has a very low acceptance rate, i.e., many restarts are required to build any new allowed configuration.

We compare the results of the hierarchical and naïve algorithms in Figure S2. For chains with different lengths of 8, 14, or 26 residues, we grew 10 000 chains each with the two algorithms. The end-to-end distance distributions obtained in this way are indistinguishable, supporting the theoretical arguments for the hierarchical algorithm introduced in the main text.

### Consistency Checks for Hierarchical Chain Growth

To test the hierarchical algorithm for consistency, we grew short chains of 50 amino acids using different procedures. We explored the effects of varying (1) the fragment length, (2) the alignment and stitching region (Figure S3), and (3) the residue overlap between the fragments.

**Figure S2:**
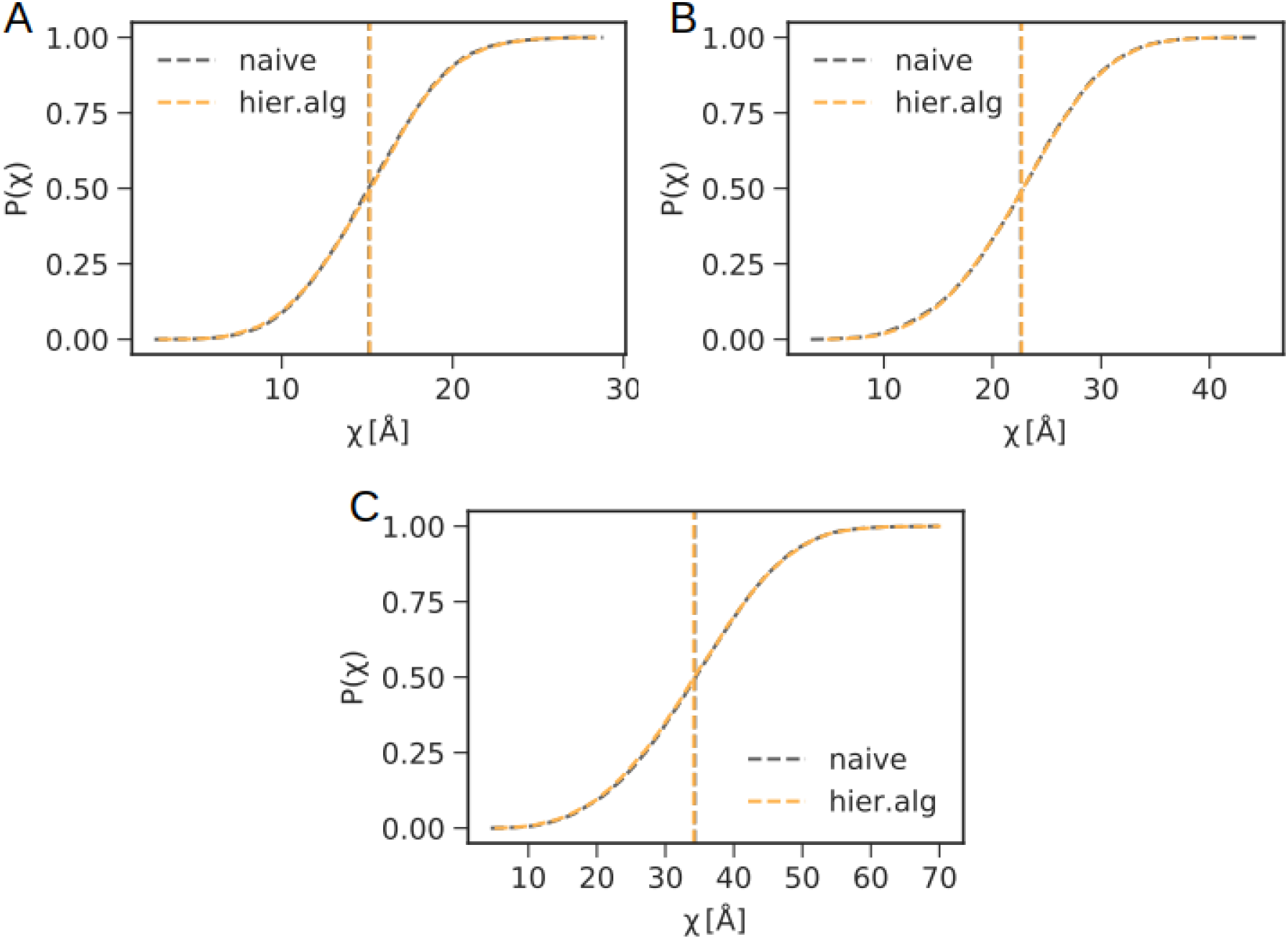
Cumulative distributions of the end-to-end distances *χ* for chains of different length grown with the naïve algorithm (blue) and the hierarchical algorithm (orange). Shown are the distributions of *χ* for chains with (A) 8 residues, (B) 14 residues, and (C) 26 residues grown with the the naïve algorithm (blue) and the hierarchical algorithm (orange).

**Figure S3:**
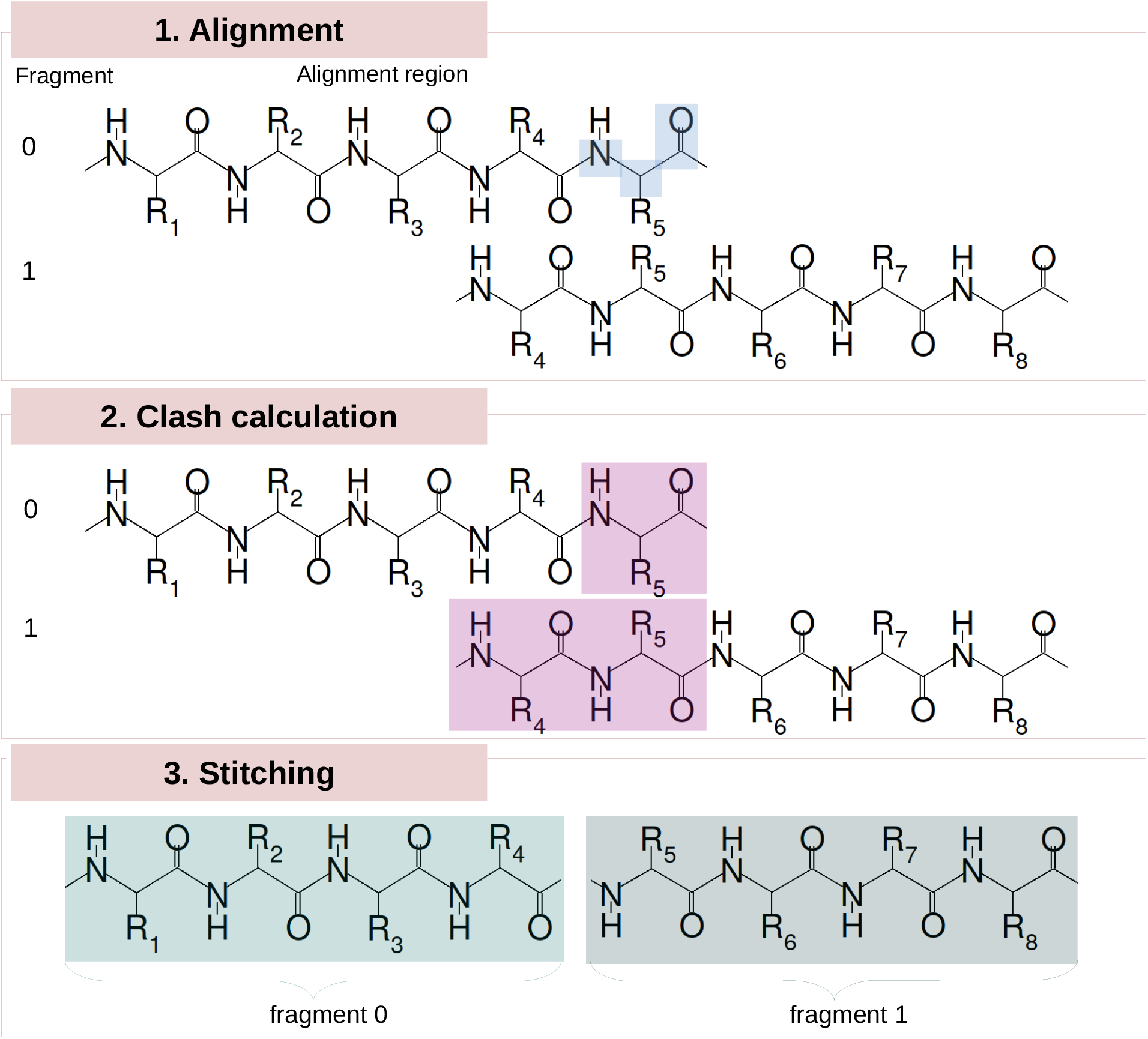
Implementation of the hierarchical Monte Carlo chain-growth algorithm with alignment of the backbone atoms. The growth procedure is shown for hierarchy level 0. To combine two randomly picked fragments, (1) the backbone atoms of the first overlapping residue in the adjacent fragments are aligned (blue shading). If the RMSD of the superimposed atoms is below a cut-off (here 0.6 Å) the aligned fragments are checked for clashes. (2) Steric overlap is probed with a cut-off distance of 2.0 Å. Atoms immediately before and after the alignment point and hydrogen atoms are excluded from the clash calculation (magenta shading). (3) If no steric clash is detected residues 1-4 from fragment 0 and residues 5-8 from fragment 1 are stored for use in the next hierarchy level.

### Fragment Length

As shown in Figure S4, the conformation of the hierarchical chain models depends somewhat on the choice of the length of the fragments from which the models are grown. Chains grown from shorter fragments tend to be more extended according to the distributions of the radius of gyration *R_G_* and of the end-to-end distance *χ*. This dependence is as a consequence of the simplifying assumption of considering only sterics in chain growth from fragments. As a result, favorable inter-fragment interactions are not accounted for, which could favor more compact structures.

### Alignment and Stitching Procedure

As shown in Figure 4A-D of the main text, the models grown by the hierarchical chain-growth approach are somewhat dependent on the alignment procedure and the stitching region chosen to merge fragments together. Aligning the backbone atoms around *C*_α_ atoms via rigid body superposition, as illustrated in Figure S3, does lead to a disruption of the dihedral angle distribution right at the stitching site (compare Figure 4A and B in the main text). This in turn leads to a bias against α-helical conformations (*ψ* < 0). This bias is largely removed by merging the fragments at the peptide bond and performing the alignment and stitching as shown in Figure 3B in the main text. As shown in Figure S4, alignment of the peptide bond also results in a slight improvement of the fragment-length dependence of *R_G_* and *χ* distributions.

### Fragment Overlap

Aligning the peptide bond between the residues present in adjacent fragments as shown in Figure 3B in the main text, we tested whether the extent of the residue overlap between merged fragments influences the overall properties of the assembled models. Figure S5 shows only small differences between an overlap of one or two residues in the radius of gyration (A), the end-to-end distance (B), and the pairwise RMSD (C) calculated from ensembles consisting of 10 000 models for a chain with 50 amino acids.

**Figure S4:**
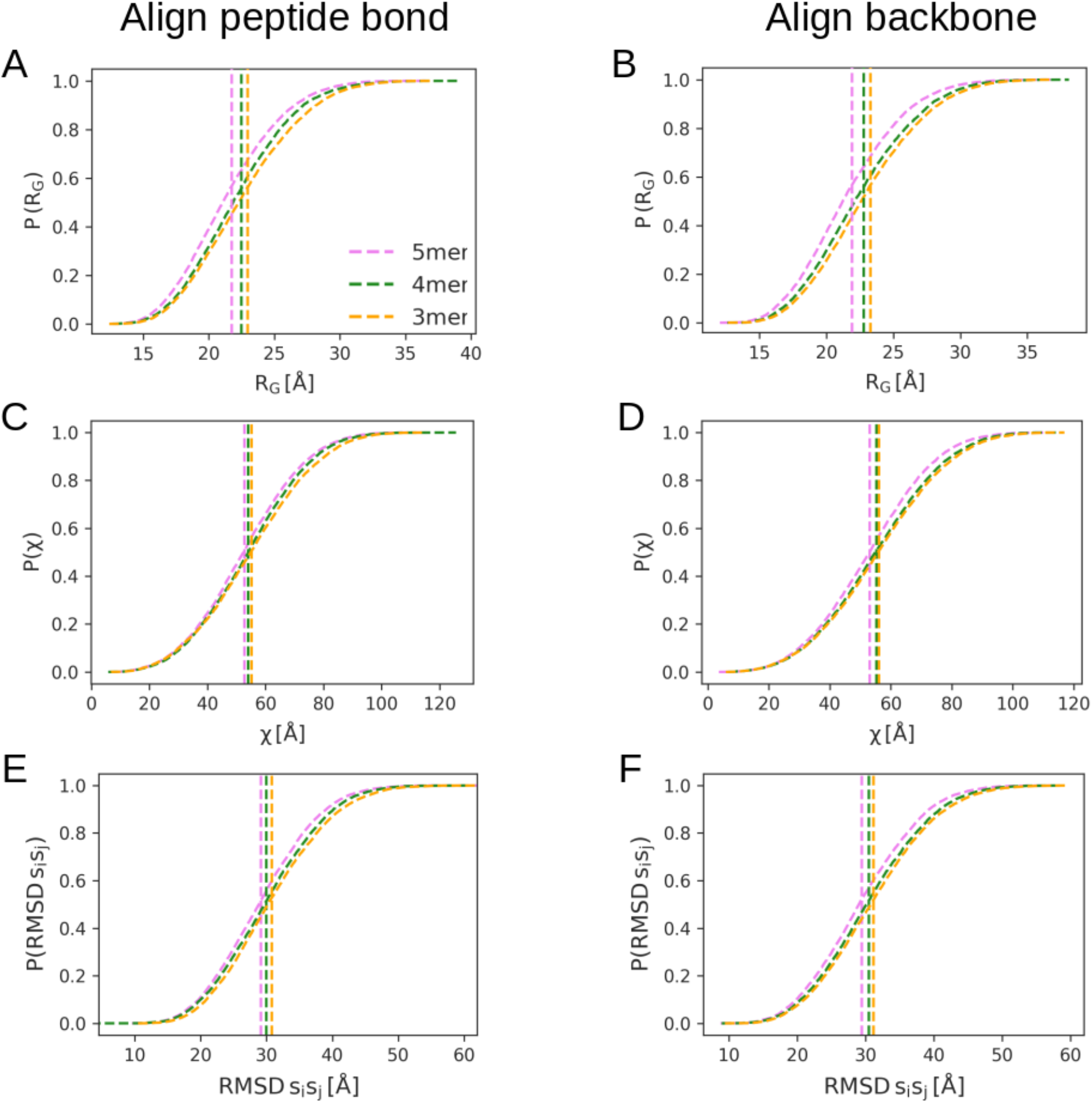
Comparison of different alignment procedures for models grown from fragments of different length. Cumulative distributions of the radius of gyration (A, B), end-to-end distances (here *χ*; C and D) and pairwise RMSD (E, F) of 10 000 models of a chain with 50 amino acids, each grown from pentamer (pink), tetramer (green), and trimer fragments (orange). Panels on the left and on the right show the distributions for models grown via alignment of the peptide bond and via alignment of the backbone atoms around *C_α_*, respectively. Vertical dashed lines indicated the respective mean values.

**Figure S5:**
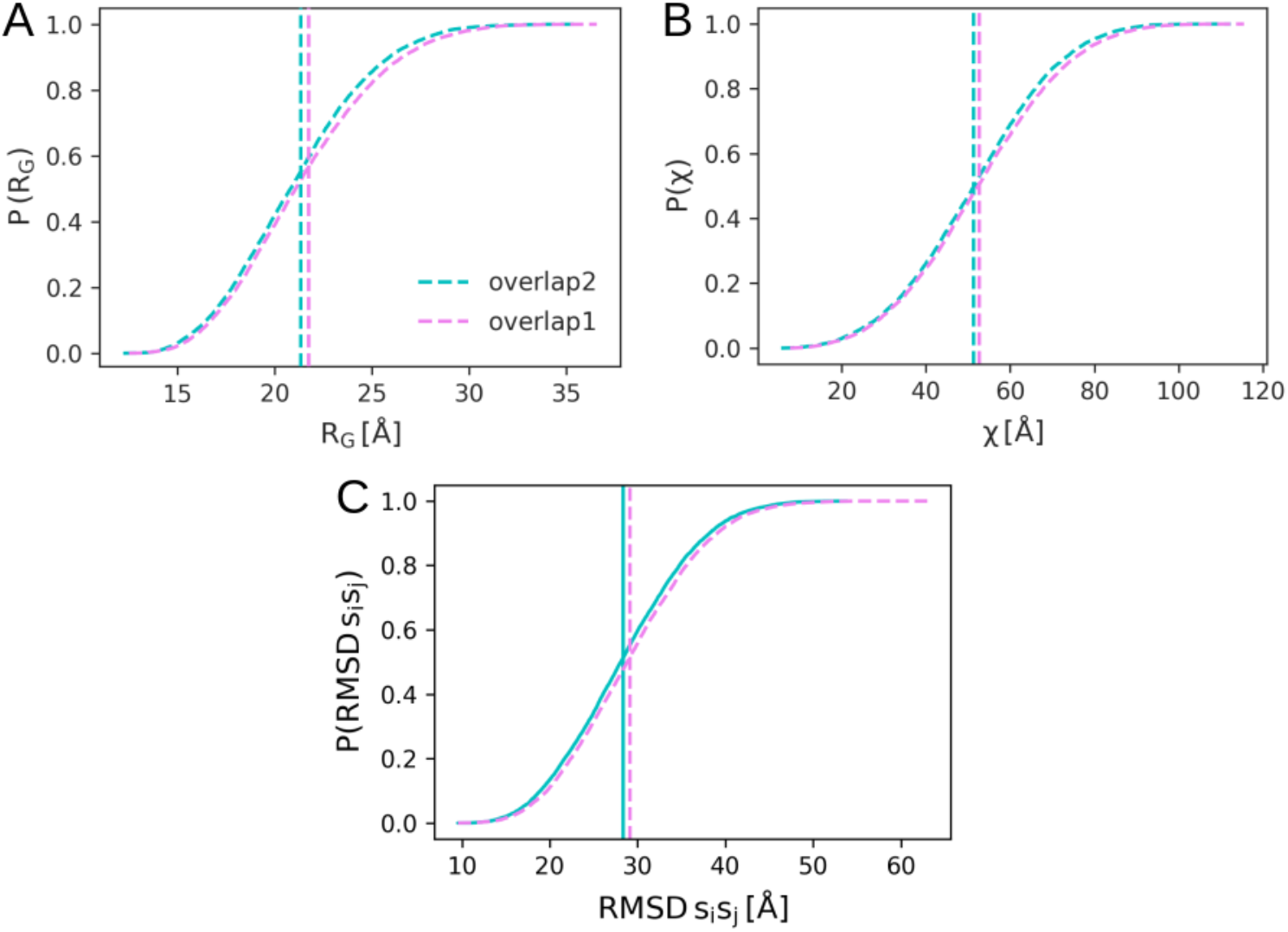
Effect of residue overlap between pentamer fragments on the structure of assembled chains. Cumulative distributions of the radius of gyration (A), the end-to-end distance (B), and the pairwise RMSD (C) of 10 000 models of a chain with 50 amino acids, each grown from pentamers with either 1 or 2 overlapping residues in adjacent fragments. Vertical dashed lines indicated the respective mean values.

### Chemical Shifts for Fragments and Full-Length Models

Assembling full-length chains by aligning on the rigid peptide bond rather than the more flexible backbone at the merge point improves the NMR chemical shifts, as predicted for the MD fragments and the full-length model after the assembly. Figure S6A-D demonstrates that by growing the full-length models via backbone alignment (right column) a bias is introduced, shifting the Δppm chemical shifts predicted for the models towards more negative values relative to the fragment values (compare main text Figure 4C and D). By contrast, when growing full-length chains via alignment on the rigid peptide bond, Δppm chemical shifts of the assembled models agree well with those of the MD fragments (Figure S6 left column).

**Figure S6:**
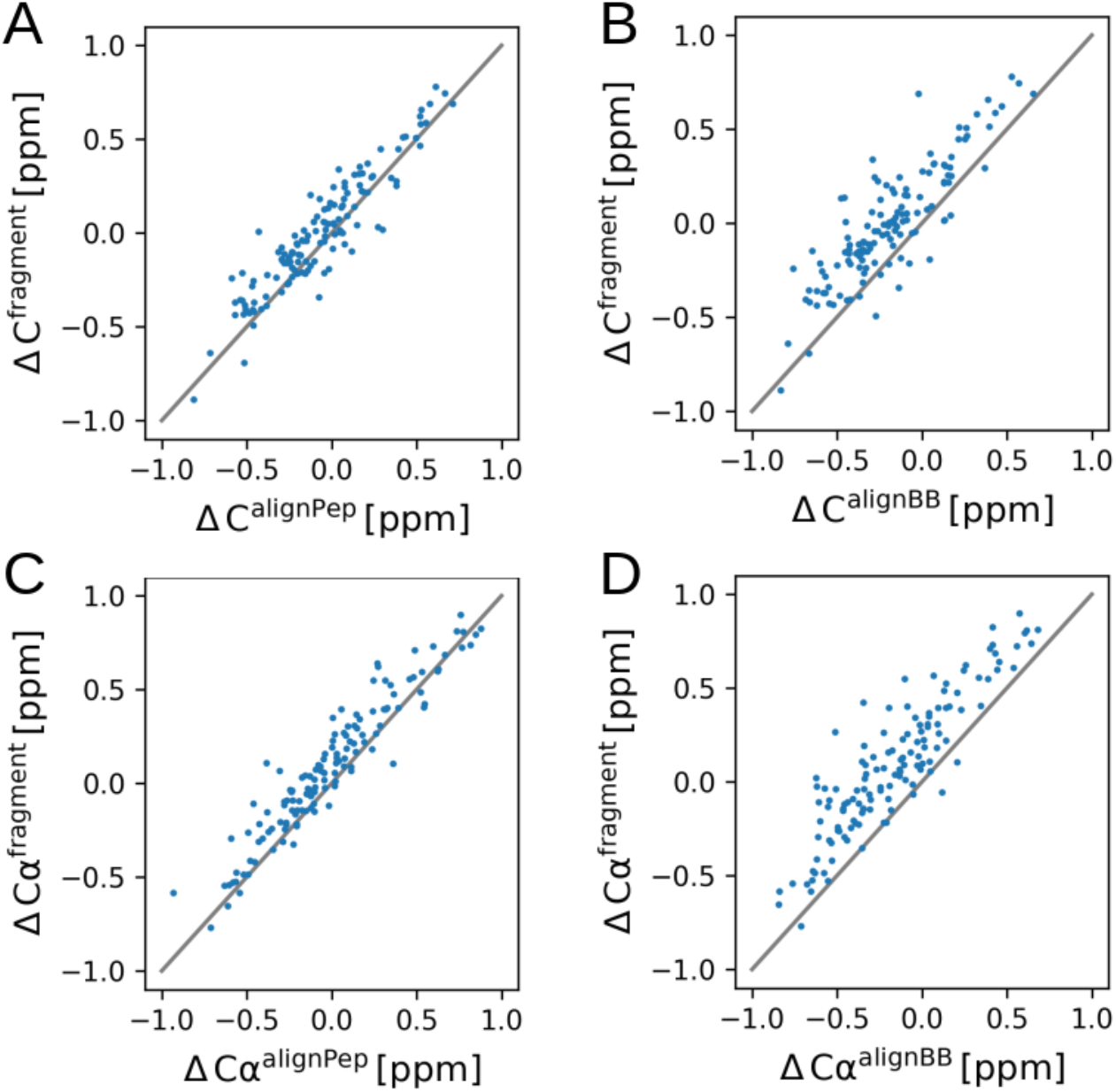
Correlation of secondary chemical shifts predicted for MD fragments and 10 000 full-length models. Left column: models grown via alignment of the peptide bond. Right column: models grown via alignment of the backbone.

